# Head-to-tail peptide cyclization: new directions and application to urotensin II and Nrf2

**DOI:** 10.1101/2022.01.05.475045

**Authors:** Yasaman Karami, Samuel Murail, Julien Giribaldi, Benjamin Lefranc, Florian Defontaine, Olivier Lesouhaitier, Jérôme Leprince, Sjoerd J. de Vries, Pierre Tufféry

**Affiliations:** Université Paris Cité, CNRS UMR 8251, INSERM U1133, Paris, France; Institut des Biomolécules Max Mousseron, UMR 5247, Université de Montpellier-CNRS, Montpellier, France; Université de Rouen Normandie, INSERM U1239 NorDiC, Neuroendocrine, Endocrine and Germinal Differentiation and Communication, INSERM US51 HeRacLeS, Rouen, France; Université de Rouen Normandie, UR CBSA, Research Unit Bacterial Communication and Anti-infectious Strategies, Evreux, France

## Abstract

Backbone head-to-tail cyclization is one effective strategy to stabilize the conformation of bioactive peptides, preventing enzymatic degradation and improving their bioavailibility. However, very little is known about the requirements to rationally design linkers for the cyclization of linear peptides. Recently, we have shown that large scale data-mining of protein structures can lead to the precise identification of protein loop conformations. Here, we transpose this approach to head-to-tail peptide cyclization. We first show that given a linker sequence and the conformation of the linear peptide, it is possible to accurately predict the cyclized peptide conformation improving by over 1 Å over pre-existing protocols. Secondly, and more importantly, we show that it is possible to elaborate on the information inferred from protein structures to propose effective candidate linker sequences constrained by length and amino acid composition. As experimental validation, we first apply our approach to design linkers for the head-to-tail cyclizations of a peptide derived from Nrf2. The designed cyclized peptide shows a 26-fold increase in binding affinity. We then consider urotensin II, a cyclic peptide already stabilized by a disulfide bond, that exerts a broad array of biological activities. The designed head-to-tail cyclized peptide, the first synthesized bicyclic 14-residue long urotensin II analogue shows an excellent retention of in vitro activity. Overall, we propose the first framework for the rational peptide head-to-tail cyclization and reveal its potential for cyclic peptide-based drug design.

## Introduction

Several naturally occurring cyclic peptides constitute alternatives to antibiotics, and peptide backbone cyclization is frequently used in peptide-based drug design to convey druggable properties to linear bioactive sequences [1, 2]. Peptides in general combine high affinity with high target selectivity and low toxicity, and are a natural choice in the targeting of protein-protein interactions. While preserving these favorable properties, peptide cyclization additionally confers peptides with more rigid conformation and enhanced stability towards enzymatic proteolysis and improves the permeability through biological barriers [3–9]. Moreover, many natural-occurring cyclic peptides are known from different kingdoms of organisms, exhibiting diverse biological activities, including anti-tumor [10, 11], antimicrobial [12, 13] and anthelmintic activities [14–16]. Together, this has caused a growing interest toward cyclic peptides, thus the number of designed cyclic peptide drugs is growing [17, 18]. However, the design of cyclic peptides remains challenging, and particularly the compatibility with a desired structure [9, 19, 20].

When designing new cyclic peptides, there are broadly two strategies that can be followed: *i) de novo* design, or *ii)* cyclization of an existing peptide. For the first strategy, a number of experimental techniques are available, such as SICLOPPS [21], phage display, and mRNA display [22]. These are all based on libraries of random cyclic peptides that are subjected to *in vitro* selection. They can be complemented with library-based computational approaches such as from Slough et al. [23], CAESAR [24], Omega [25] and CycloPs [26] based on rdkit (https://github.com/rdkit/rdkit). Those approaches are conceptually similar to the molecular modeling of small ligands, with the corresponding strengths (arbitrary molecular topologies) and weaknesses (limited number of flexible bonds). For computational *de novo design*, some recent approaches focusing on protein-protein interactions use hot spots identified at the interface to identify linker residues or template cyclic conformations matching the geometry of the hot spots [27, 28]. They have not been experimentally validated so far. An alternative approach is to perform peptide structure prediction, using one of the many fragment-based methods that are available, such as PLOP [29, 30], Peplook [31, 32], PEPstrMOD [33] or PEP-FOLD [34, 35], while imposing cyclization as a bond or distance restraint (see [36] for a review). Since these methods leverage the existing wealth of knowledge of protein and peptide structure, they can deal with larger peptides, but have difficulties where this knowledge falls short, *i*.*e*., for unnatural amino acids.

For the second strategy, the starting point is an existing linear peptide of known structure. It is well established that small linear peptides generally exist in solution in an interchangeable conformational equilibrium. This flexibility provides to bioactive peptides the ability to interact with several types or subtypes of receptors for instance. Stabilizing a bioactive conformation is a challenge that can be tackled by a variety of cyclization strategies. On the one hand, this can be as straightforward as mutating two spatially close residues into cysteins with the aim of introducing a disulfide bond. On the other hand, sophisticated chemical scaffolds or cyclotides can be used for the grafting or stitching of peptides or cyclotides into rigid bioactive conformations [37, 38]. One particular successful strategy has been head-to-tail peptide backbone cyclization [39–48]. This involves the design of a sequence that links the N- and C-terminal extremities of the peptide. In principle, any amino acid can be part of the linker sequence, but Gly, Ala and Pro residues are often favored because they are small and their side chains cannot form hydrogen bonds, which could potentially disrupt the bioactive conformation.

Head-to-tail cyclization leads to cyclic peptides with improved pharmacological properties (affinity, potency, efficiency, selectivity) when compatible with target specificity (or bioactivity conservation). Whether the cyclic peptide is active or not, it is generally less sensitive to metabolic degradation. However, cyclization is often unsuccessful due to imposed conformational restriction that is too strict and too far from the bioactive structure. In order to avoid this, it is necessary to understand the general sequence-structure requirements; in particular: what is the allowed sequence space of the linker, and what will be the structure of the cyclized peptide? This is a challenging issue, and to the best of our knowledge, there is only one computational protocol that has been successfully applied to head-to-tail cyclization linker design, namely the Rosetta protocol used by Bhardwaj et al. [6]. However, in that study, the sequence and structure of the entire cyclic peptide were designed from scratch. Otherwise, we are not aware of any computational methods that can predict the sequence and structure of a head-to-tail cyclization linker, while preserving the sequence and structure of the linear peptide that is being cyclized.

Recently, we have developed a fast data-based approach that identifies loop candidates mining the complete set of available experimental protein structures [49, 50]. This is done by treating the loop as a gap in the structure, and considering the flanking regions of the structure immediately before and after the gap. Loop candidates are then favored that *i)* superimpose well onto the flanks, and *ii)* have a compatible sequence. Recognizing the conceptual similarity, we have now developed a method that extends this approach and applies it to rational head-to-tail peptide cyclization. It addresses two complementary concerns: *i)* given a sequence for the cyclization linker, it can predict structural models for the cyclized peptide, *ii)* it can propose candidate cyclization linker sequences, constrained by length and amino acid composition. It is the first method that can propose the sequence or the structure of a headto-tail cyclized peptide, starting from the linear peptide structure. For structure prediction, it was validated on a benchmark of five cyclic conotoxin structures for which a linear structure is available as well. With regard to the experimental structures, the predicted cyclized peptide models had a root-mean-square deviation (RMSD) of 2.0 Å (3.2 Å) for the top 20 (top 1) models, an improvement of more than 1 Å over the Rosetta Next-generation KIC (NGK) protocol [51], a high-resolution Rosetta protocol for the modelling of missing regions. For sequence prediction, it was validated on the same benchmark and in result, experimental sequences were ranked significantly better than other sequences of the same length and composition.

As a functional validation, this approach was used to design head-to-tail cyclized variants of two peptides. The first one corresponds to the interaction of a 9-residue long fragment of Nrf2 in interaction with Keap1, as previously crystallized in linear form [52]. The second one corresponds to the human urotensin II (UII), that is an 11-residue long disulfide-bridged peptide [53]. UII exerts a broad array of biological activities, in particular in the central nervous system, the cardiovascular system, and the kidney. It has been suggested that the cognate receptor of UII (UT), may emerge as a valuable and innovative therapeutic or diagnostic target [54]. Indeed, high affinity, potent and selective UT peptide ligands have been designed, from structure-activity relationship studies [55] to further elucidate the pharmacology and biology of UII towards new therapeutic opportunities, such as the treatment of sepsis-induced lung damage [56]. In this context, mastering the introduction of a main conformational restraint through head-to-tail cyclization could become a standard strategy to improve pharmacological profile of peptide ligands [57].

## Results

Our approach (PEP-Cyclizer) considers all cyclization linker candidate structures that are compatible with the flanks of the uncyclized peptide structure. The sequences of these linker candidates, potentially filtered by *a priori* sequence constraints, are then used to build a linker sequence profile. This profile feeds a Hidden Markov Model from which it is possible to estimate the likelihood of candidate linkers using a forward-backtrack algorithm. Alternatively, if the linker candidates are restricted to one known linker sequence, they are clustered and superimposed onto the flanks, providing structural models of the cyclized peptide. **Figure 1** depicts the workflow of the method.

**Figure 1:**
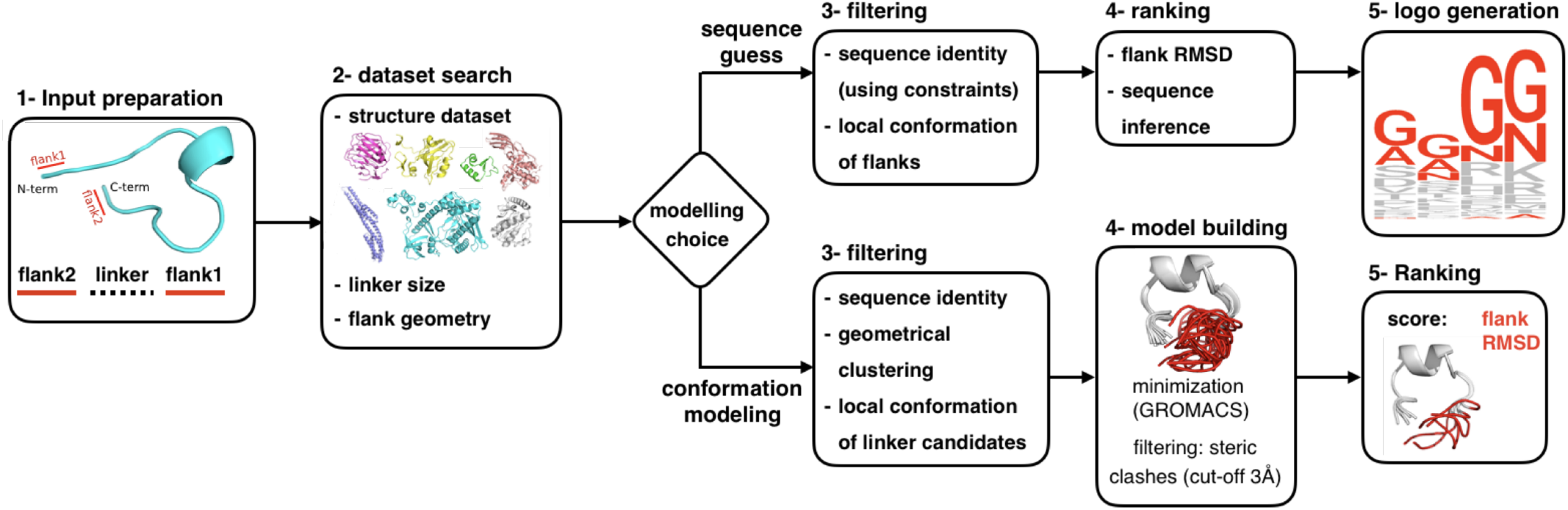
The workflow of PEP-Cyclizer. The workflow describes main steps for peptide head-to-tail cyclization. The method provides two possibilities: proposing candidate sequences for the linker, or modelling the 3D conformation. The steps of the workflow are: input preparation, linker candidate search, candidate filtering, model building, model selection and logo generation in case of sequence prediction. The inputs are a linear peptide and either the amino acid constraints for sequence prediction, or the linker sequence for conformation modelling. In the final step, for conformation modelling, the 20 best models are returned as the final predictions. For sequence prediction a logo is generated and a forward-backtrack algorithm is used to sample the sequence space and assess the likelihood of the candidate linkers. Note that the sequence logo serves strictly as a global visualization of the ensemble of generated sequence candidates, and has no predictive power by itself.

As a positive control, PEP-Cyclizer was applied to 64 cyclic peptides from the CyBase database (http://www.cybase.org.au/) [58, 59] (see **Table S1** for a complete list of studied peptides). 1147 linear peptides were artificially generated by removing segments of 2-7 residues from the 64 cyclic peptides, details are reported in **Supplementary Materials - CyBase benchmark**. Unlike a real-world situation, where a peptide may undergo conformational changes upon cyclization, these artificial linear peptides represent perfect conformations for modelling the removed linker conformation. For all linker sizes, our protocol was able to produce accurate models of the local linker conformation, with an average accuracy of 1 Å or better. This is comparable (although not fully equal in accuracy) to models obtained for the same peptides using Rosetta NGK (comparisons reported per peptide and linker size in **Table S2** and **S3, respectively**).

The protocol was then applied to a small benchmark of real-world cases, in the form of several conotoxin peptides where both cyclized and non-cyclized structures are available in the PDB. Seven distinct cyclized/non-cyclized pairs of peptide structures were identified (**Table 4** and **S4**). The range of backbone RMSD between the overlapping region of cyclized and uncyclized forms is 0.4-2.5Å. Using the known linker sequence, only the non-cyclized structure was used to model the linker. In this case, PEP-Cyclizer was able to return a model approximating the global structure of the linker at 2.01Å on average (1.01Å for the local linker conformation), as reported in **Table 1**. This is a considerable improvement over Rosetta NGK (3.48Å), which suffers from the structural imprecisions caused by conformational change upon cyclization. In contrast, our results show PEP-Cyclizer to be rather robust against such imprecisions. **Figure 2** illustrates the results for the best predictions - out of the top 20 - of PEP-Cyclizer (in green) and Rosetta NGK (in cyan), starting from the first NMR conformation of each uncyclized peptide.

**Table 1:** Comparison of RMSD values for the predicted linkers using all the NMR models of the linear peptides. For each peptide we report the average local and global RMSD values over the top 1 (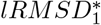 and 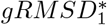) and best out of top 20 predictions (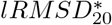 and 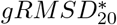). The average values are measured over all *N*_*uncyclized*_ NMR conformations of each linear peptide (see **Methods**). The RMSD values are calculated over the backbone atoms (N, C, C*α* and O). ^+^The structure of 1mii and 1m2c correspond to the same protein (*α*-Conotoxin MII), and to avoid redundancies in reported values, we measured the average considering the best predictions between 1m2c and 1mii for each method.

| Name | $l_{sz}$ | $N_{model}$ | PEP-Cyclizer | | | | Rosetta NGK | | | |
| --- | --- | --- | --- | --- | --- | --- | --- | --- | --- | --- |
| | | | $lRMSD_{20}^*$ | $gRMSD_{20}^*$ | $lRMSD_1^*$ | $gRMSD_1^*$ | $lRMSD_{20}^*$ | $gRMSD_{20}^*$ | $lRMSD_1^*$ | $gRMSD_1^*$ |
| 2ew4 | 2 | 20 | 0.47±0.13 | 2.03±0.84 | 0.67±0.18 | 3.57±1.35 | 0.31±0.22 | 2.53±1.25 | 0.39±0.26 | 3.10±1.25 |
| 1ixt | 3 | 20 | 0.46±0.08 | 2.31±0.25 | 1.43±0.35 | 3.52±1.50 | 0.37±0.14 | 2.81±0.54 | 0.43±0.14 | 3.08±0.66 |
| 1m2c | 6 | 14 | 1.31±0.15 | 1.99±0.16 | 1.66±0.12 | 2.58±0.57 | 1.33±0.17 | 4.58±0.90 | 1.53±0.16 | 6.08±1.53 |
| 1mii<sup>+</sup> | 6 | 20 | 1.35±0.01 | 1.72±0.01 | 1.75±0.50 | 3.11±1.74 | 1.56±0.11 | 5.76±0.55 | 1.72±0.12 | 7.22±0.45 |
| 2h8s | 6 | 20 | 1.24±0.12 | 2.12±0.10 | 2.03±0.29 | 4.05±1.21 | 1.30±0.19 | 3.01±0.53 | 1.60±0.21 | 3.81±0.64 |
| 1m2c | 7 | 14 | 1.59±0.16 | 1.97±0.25 | 2.09±0.56 | 3.66±1.56 | 1.84±0.37 | 5.34±1.30 | 2.12±0.26 | 6.53±1.29 |
| 1mii<sup>+</sup> | 7 | 20 | 1.51±0.08 | 1.89±0.06 | 1.69±0.01 | 2.30±0.02 | 1.56±0.03 | 5.51±0.72 | 1.64±0.06 | 7.80±0.67 |
| average<sup>+</sup> |  |  | 1.01±0.46 | 2.01±0.43 | 1.49±0.52 | 3.24±1.26 | 0.95±0.63 | 3.48±1.39 | 1.13±0.71 | 4.28±1.77 |

**Figure 2:**
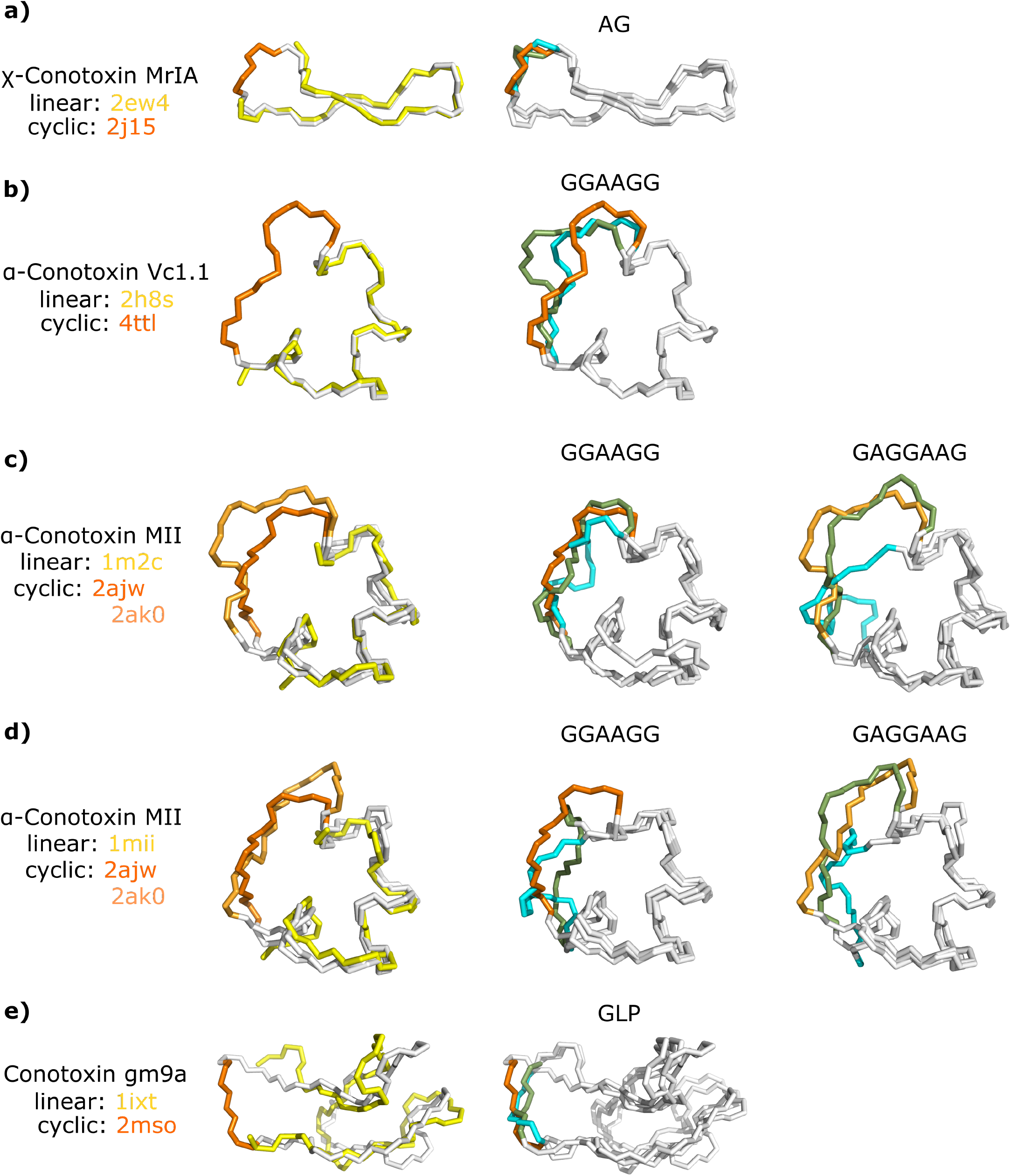
Structure of the studied linear conotoxins and their corresponding engineered cyclic peptides. The native linear and cyclic peptides are shown at the left column, colored in yellow and orange, respectively. The structures on the middle and right columns, represent the comparison between the native linker (in orange), linkers modelled by PEP-Cyclizer (in green) and Rosetta NGK (in cyan). See **Table 1** for the corresponding *gRMSD*_20_ (and *lRMSD*_20_) values. The corresponding linker sequences are reported for every model, at the top.

Next, the ability of our approach to propose peptide linker sequences was tested. The same conotoxin benchmark was used, adding ten cyclic sequences with available structures for the uncyclized but not the cyclized peptide, for a total of seventeen sequences. The details of the peptides are reported in **Tables 4** and **S4**. As potential linker sequences, all combinations of all amino acids that are present in the experimental linker sequence (typically only Gly and Ala) were considered, and ranked by the forward-backtrack algorithm. The results are shown in **Figure 3**. The experimental sequences were ranked significantly better (average percentile: 37.4, p=0.025) than other potential sequences.

**Figure 3:**
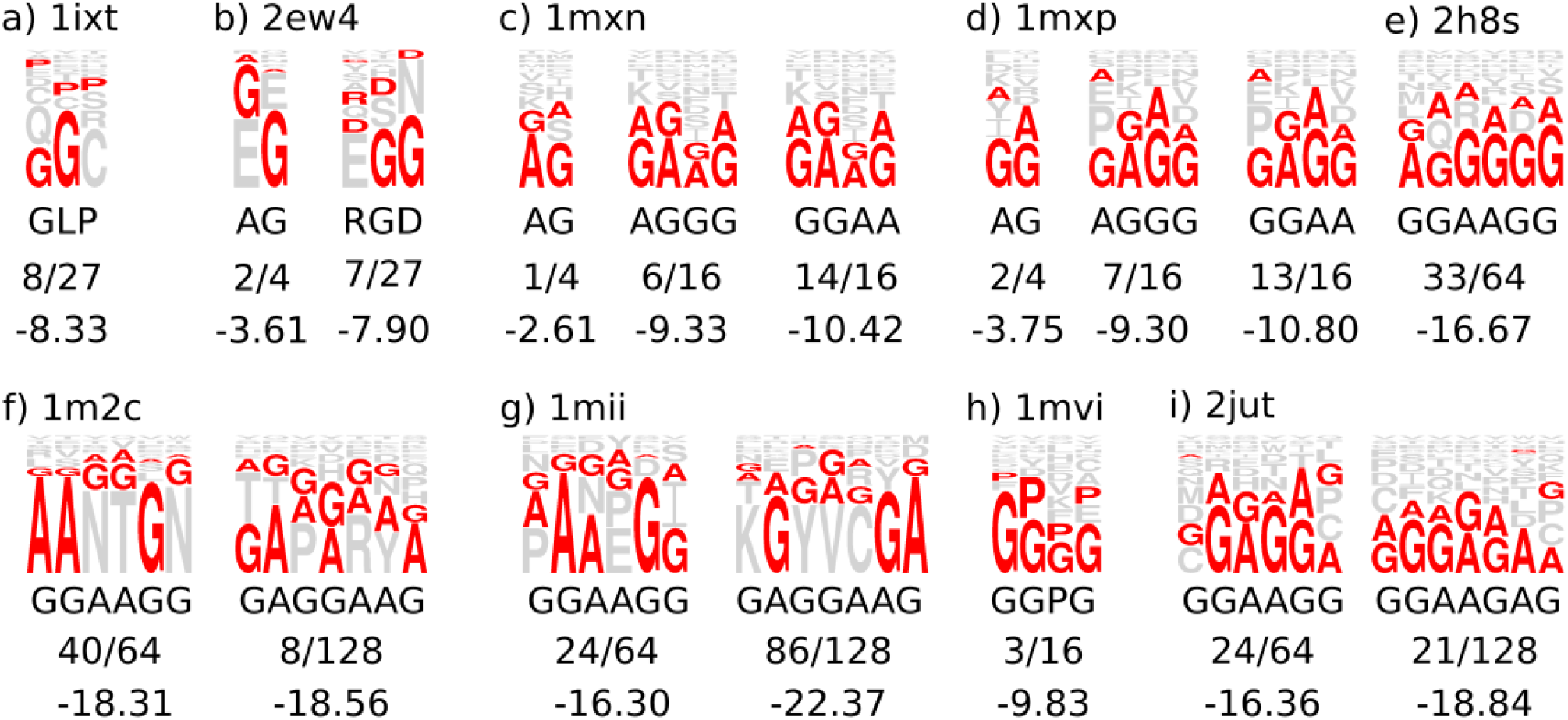
Sequence logo generated by PEP-Cyclizer for the studied cases. The pdb code of the linear peptides used as input are reported for each case. Below every logo, the desired linker sequence, its rank and score among the proposed sequences by the forward-backtrack algorithm are reported.

### Application to the design of cyclized Nrf2 peptides

We first applied our head-to-tail cyclization protocol to design linkers to cyclize the peptide corresponding to residues 76-85 of the Nrf2 protein. This peptide has been crystallized in interaction with the Keap1 protein [52]. The peptide conformation of the crystal structure was used as starting point. For cyclization, a linker of size 3 was considered, accepting only alanine, proline and glycines. The restriction to these amino types alone results from the concern to limit interference with the binding mode of the peptide as much as possible, by favoring the amino acids known to have a rather neutral behavior, as done in other studies []. **Table 2** shows all 27 possible linkers ranked by decreasing likelihood. Due to the difficulty to synthesize such cyclic peptides, we focused our efforts on two different linkers, namely the highly ranked AGG and the poorly ranked PAA.

**Table 2:** The likelihood of each of the possible 27 linker sequences for Nrf2.

| linker | L |
| --- | --- |
| GGG | -5.71 |
| AGG | -5.73 |
| GAG | -6.32 |
| PGG | -6.71 |
| APG | -7.56 |
| AAG | -7.65 |
| GPG | -7.79 |
| PAG | -7.84 |
| PPG | -8.34 |
| GGA | -8.71 |
| AGA | -8.74 |
| GGP | -9.66 |
| AGP | -9.68 |
| PGA | -9.71 |
| GAP | -10.01 |
| GAA | -10.63 |
| PGP | -10.66 |
| APP | -11.06 |
| APA | -11.09 |
| GPP | -11.30 |
| GPA | -11.32 |
| PPP | -11.85 |
| PPA | -11.87 |
| AAA | -11.97 |
| PAA | -12.16 |

Three Nrf2-derived peptides were synthesized (see Methods), namely the original linear *Nrf* 2_(76*−*85)_ peptide and the same peptide cyclized with the two linkers. The sequences of the peptides are shown in **Figure 4**.

**Figure 4:**
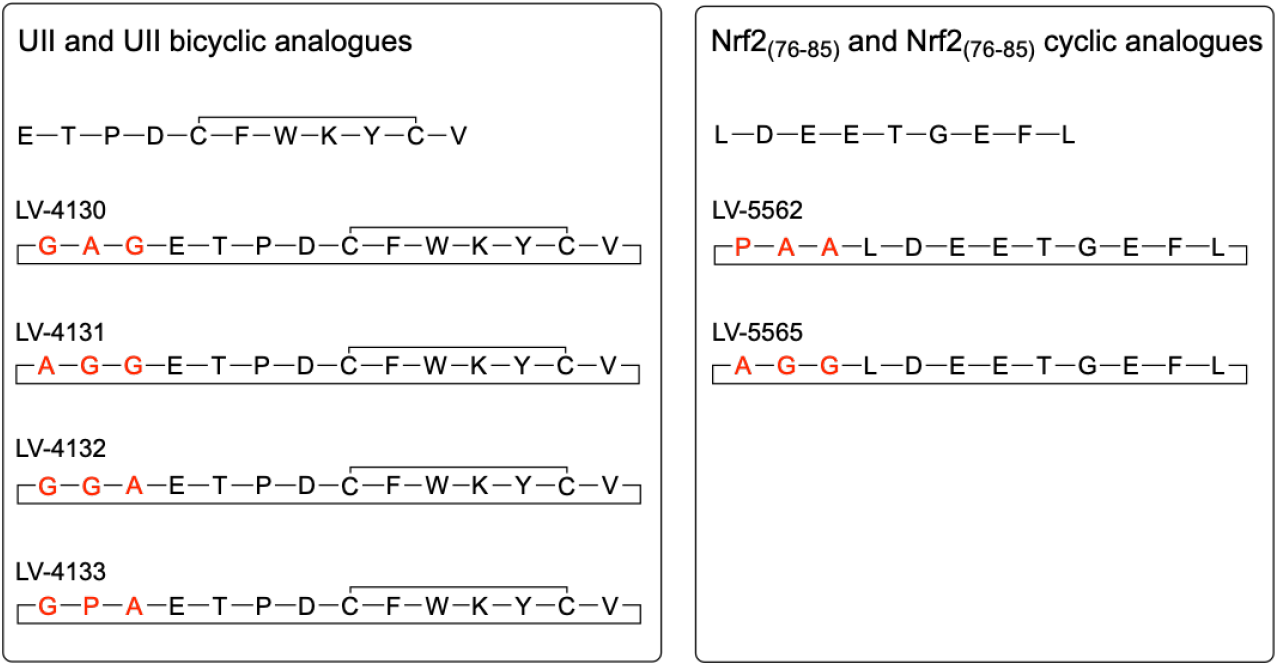
Sequence of human UII, human/mouse *Nrf* 2_(76_*−* _85)_ and head-to-tail cyclic analogues. Linker residues for ring closing are marked in red.

Using Microscale Thermophoresis (MST), we then measured the binding affinities between his-tagged Keap1 protein and each of the synthesized peptides. The linear Nrf2 peptide (LV-5554) possessed an affinity for Keap1 with a Kd measured at 3.16 *±* 0.64*µM* **Figure 5**. Under the same conditions, the peptide LV-5565 (linker AGG) possessed a stronger affinity for the Keap1 protein with a Kd value of 0.12 *±* 0.04*µM*, 26 times better than that measured for linear Nrf2. Finally, the other cyclic peptide LV-5562 (linker sequence PAA) possessed an affinity slightly better than the linear peptide (Kd = 1.92 *±* 0.44*µM*). Overall, the highly ranked linker indeed showed a much higher affinity than the linker that is ranked last, although the latter still resulted in a cyclized peptide that had a binding affinity comparable to the linear peptide.

**Figure 5:**
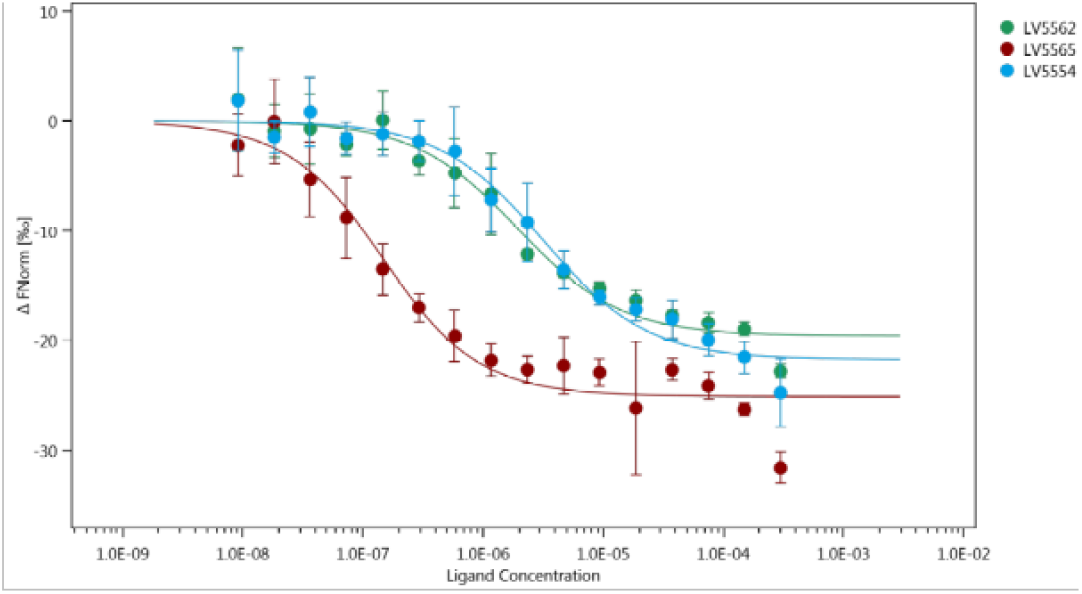
Dose response fit of MST for cyclized Nrf2 peptides. MST dose response curves for interaction between Keap1 labelled protein and the linear *Nrf* 2_(76_*−*_85)_ peptide (LV-5554, blue circles), the LV-5562 peptide (green circles) and the LV-5565 peptide (red circles). The data are the mean of three independent experiments for both cyclic (LV-5562 and LV-5565) peptides and of five independent experiments for the linear *Nrf* 2_(76_ *−*_85)_ peptide (LV-5554). The thick lines correspond to the fit from the law of mass action. (Fl) fluorescently labelled; (FNorm) normalized fluorescence. ΔFnorm values of Y-axis in the dose response plot are calculated from the ratio of normalized fluorescence F0/F1, where F0 corresponds to the normalized fluorescence prior to MST activation. F1 is determined after an optimal MST power-dependent time interval which yields the best signal-to-noise ratio.

### Application to urotensin II

We then turned to the prediction of a linker sequence for the head-to-tail cyclization of UII. So far, only the structures of a fragment corresponding to the eight last amino acids of UII and its N-methylated tryptophan counterpart, [(N- Me)Trp^7^]U-II_4*−*11_ in polar conditions (PDB entries 6HVB and 6HVC) have been solved by NMR. Since our goal was to obtain a head-to-tail cyclized version of UII, we decided to start from 3D models of the complete UII (11 amino acids) which includes one disulfide bond. Therefore two ensembles of 8 and 5 conformations were generated using two distinct strategies: *i)* molecular dynamics simulations (MD) and *ii)* PEP-FOLD [35]. The models are highly structurally divergent (**Supplementary Figure S2**), with typical RMSD values in excess of 2 Å both within and between the ensembles (**Supplementary Table S6**). Consequently all those models were used as the starting points for the cyclization (see **Methods**). Based on the average distance between the N- and C-terminus of the models (7.27 *±* 2.14 Å), a linker of size 3 was considered, accepting only alanine, proline and glycines. **Table 3** presents the results accumulated for each of the two ensembles of models. As can be observed, it is striking that despite the diversity of the conformations and the way they were obtained, those two independent ensembles of models resulted in a rather stable ranking of the predicted sequences. This is reflected by the fact that in both cases, the top 4 consists of the same four sequences, as well as by the high overall correlation of the ranks (Spearman r=0.98).

**Table 3:** The likelihood of each of the possible 27 linker sequences. Two independent series of models (8 generated using MD and 5 using PEP-FOLD) were used as starting points.

| PEP-FOLD |  | MD |  |
| --- | --- | --- | --- |
| linker | L | linker | L |
| AGG | -7.42 | AGG | -7.21 |
| APG | -7.65 | GAG | -7.52 |
| GAG | -7.67 | PGG | -7.52 |
| PGG | -7.73 | APG | -7.61 |
| AGA | -7.80 | GGG | -7.64 |
| PAG | -7.81 | PAG | -7.69 |
| GGG | -7.89 | AAG | -7.78 |
| AAG | -7.90 | AGA | -7.80 |
| PGA | -8.11 | PGA | -8.10 |
| GAP | -8.12 | PPG | -8.11 |
| PPG | -8.15 | GAP | -8.21 |
| AGP | -8.19 | AGP | -8.22 |
| PAP | -8.26 | GGA | -8.23 |
| GGA | -8.27 | GPG | -8.36 |
| APP | -8.29 | PAP | -8.38 |
| APA | -8.32 | AAP | -8.47 |
| AAP | -8.34 | APA | -8.49 |
| GPG | -8.44 | APP | -8.50 |
| PGP | -8.50 | PGP | -8.53 |
| GGP | -8.66 | GGP | -8.65 |
| GAA | -8.75 | GAA | -8.79 |
| PPP | -8.80 | PAA | -8.96 |
| PPA | -8.82 | PPA | -8.99 |
| PAA | -8.88 | PPP | -9.00 |
| AAA | -8.97 | AAA | -9.06 |
| GPP | -9.08 | GPA | -9.24 |
| GPA | -9.11 | GPP | -9.25 |

**Table 4:** The list of real cases for head-to-tail cyclization. The PDB code of the un-cyclized and cyclized peptides (if available) are reported. With the exception of 4ttl, all the other structures are obtained using NMR and has several models. The average RMSD values are measured between all the models of the un-cyclized and cyclized conformations. In some cases more than one linker sequence exist, as reported in the last column of the table.

| un-cyclized | #NMR models | cyclized | #NMR models | RMSD (Å) | un-cyclized size | cyclized size | linker sequence |
| --- | --- | --- | --- | --- | --- | --- | --- |
| 1m2c | 14 | 2ajw | 20 | linear 1.22 +/- 0.10 | 16 | 22 | GGAAGG |
| 1m2c | 14 | 2ak0 | 20 | 1.09 +/- 0.12 | 16 | 23 | GAGGAAG |
| 1mii | 20 | 2ajw | 20 | 1.26 +/- 0.09 | 16 | 22 | GGAAGG |
| 1mii | 20 | 2ak0 | 20 | 1.03 +/- 0.12 | 16 | 23 | GAGGAAG |
| 2h8s | 20 | 4ttl | 1 | 0.40 +/- 0.00 | 16 | 22 | GGAAGG |
| 1ixt | 20 | 2mso | 20 | 2.45 +/- 0.07 | 27 | 30 | GLP |
| 2ew4 | 20 | 2j15 | 21 | 1.03 +/- 0.40 | 13 | 15 | AG |
| 2ew4 | 20 | - | - | - | 13 | 16 | RGD |
| 1mxn (1mxp) | 20 | - | - | - | 15 | 17, 19, 19 | AG, AGGG, GGAA |
| 2jut | 20 | - | - | - | 13 | 19, 20 | GGAAGG, GGAAGAG |
| 1mvi | 15 | - | - | - | 25 | 28 | GLP |

To test the significance of our approach, we analyzed the impact of top-, mid- and bottom-ranked linkers on the agonist properties of the respective bicyclic UII analogues. Therefore, UII was cyclized with the sequences AGG, GAG, GGA and GPA as displayed in **Figure 4** leading to compounds LV-4131, LV-4130, LV-4132 and LV-4133, respectively (see **Methods**). Although obtained with low yield (*<* 1%), each bicyclic analogue was highly pure. The pharmacological profile of these synthesis-challenging compounds was assessed by measuring their ability to increase intracellular calcium concentration ([Ca^2+^]i) in human UT-transfected CHO cells (Eurofins-Cerep and EuroscreenFast) as previously described [60]. As shown in **Figure 6**, UII and LV-4130 induced a dose-dependent increase in [Ca^2+^]i with EC_50_ of 0.7 and 46 *nM*, respectively. The other analogs were less potent than LV-4130 and exhibited EC_50_ values *>* 6*µM*. Noteworthy, LV-4130 is a first bicyclic UII analogue retaining a substantial ability to increase [Ca^2+^]i in UT-transfected CHO cells. Despite a shift in potency, the EC_50_ was less than 2 orders of magnitude lower, and LV-4130 is a nanomolar active UT agonist of peculiar interest. Indeed, its backbone cyclic structure may confer a lower susceptibility to metabolic degradation and a better selectivity for UT or a subset of UT’s signaling cascade that deserves to be investigated - biased agonist concept. We also emphasize that for UII, linker design started from modelled structures, not from experimental as for the Nrf2 derived peptide.

**Figure 6:**
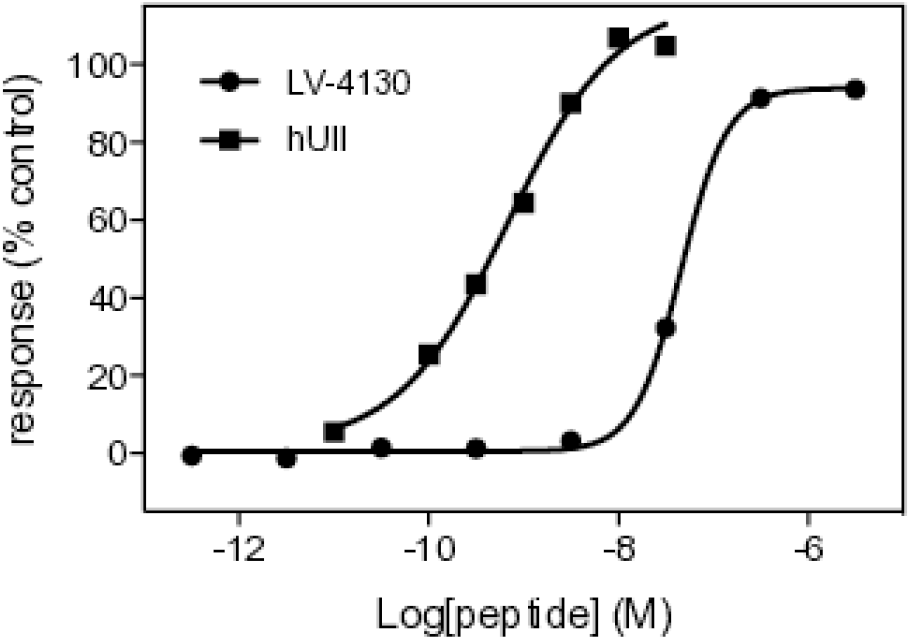
Concentration-dependent agonist-evoked Ca^2+^ responses on UT-transfected CHO cells. Agonist responses were expressed as a percent of the response observed with a maximally effective concentration of UII (100 *nM*). Data points represent mean of duplicate.

## Discussion

Recently, we have demonstrated that the current structural information available in the Protein Data Bank (PDB) [61] is now sufficient to propose accurate protein loop candidates, in a manner that is robust for conformational inaccuracies. Latest results in the field of deep learning applied to protein structure modeling largely support this observation [62]. In the present study, we asses the extension of this observation to head-to tail peptide cyclization, and introduce the first computational method to assist the design of head-to-tail cyclization of an existing peptide structure, a wellknown strategy to enhance peptide resistance to enzymatic degradation and thus peptide bioavailability. The method addresses two complementary questions, namely : *(i)* proposing candidate sequences for the linker, a facility to assist medicinal chemists, and *(ii)* predicting the 3D conformation of the linker, for further peptide conformational stability analysis or peptide-receptor docking. Up to now, there has been an evident lack of computational methods to answer those questions. Existing methods [23–26, 29–35] are oriented towards *de novo* design and do not perform head-to-tail cyclization of existing structures. We are aware of a single existing computational method, the Rosetta protocol by Bhardwaj et al. [6], that is able to design head-to-tail cyclization linkers for pre-existing peptide structures. However, in that method, what is pre-defined is the complete structure of the entire cyclic peptide, including the linker; also, the sequence of the entire peptide is designed from scratch, and not just that of the linker. In contrast, PEP-Cyclizer starts from the existing structure of a linear peptide, and predicts the sequence or structure of a cyclization linker, while leaving the rest of the peptide undisturbed. This clearly addresses a need in the field of peptide optimization and, to the best of our knowledge, it is the first computational method designed to do this.

The performance of PEP-Cyclizer to model the structure of cyclized peptides was validated on a set of conotoxins for which both linear and cyclic peptide structures are known. For comparison, we also evaluated a Rosetta protocol, not the one from Bhardwaj et al. [6], but the Rosetta NGK protocol [51], a state-of-the-art protocol for building missing loops in crystal structures. It must be mentioned that Rosetta NGK is not designed for peptide cyclization and we had to modify the input data and convert the head-to-tail cyclization to loop modelling (*i*.*e*., dividing the peptides in two segments and switching them to generate a gapped structure). In all cases, the peptide linker models generated had a significantly better global accuracy. This is especially evident for the two longest (7 amino acids) linkers, where the RMSD was *<* 2 Å, while *>* 5 Å for Rosetta NGK.

Our protocol is an extension to the problem of head-to-tail peptide cyclization of an algorithm initially designed for the problem of protein loop modeling; details about the algorithm are reported in our previous study [49]. Essentially, the linker/loop is treated as a gap in the structure, and a structural database search is performed using the flank regions on either side. Like for our previous work, our approach is a consensus method that considers both structural compatibility (*i*.*e*., good superposition of the linker candidate onto the flanks) and sequence compatibility. Therefore, when using it to predict linker conformations, it is essential to consider all 20 candidate structures. When forced to make a single prediction, the quality deteriorates considerably (from 2.0 to 3.2 Å). While a top-1 accuracy is naturally less favorable than a best-of-20 for any prediction method, it is specifically true for PEP-Cyclizer, as the effect is much weaker for Rosetta NGK (from 3.5 to 4.3 Å).

In contrast, our protocol showed very robust against conformational changes. For the conotoxin benchmark, the range of backbone RMSD between the overlapping region of cyclized and uncyclized forms is 0.4-2.5Å. This is to be compared with the positive control, where this RMSD is zero. However, the global accuracy of the PEP-Cyclizer models is essentially the same between the two (2.0 Å vs 1.87 Å). This is a stark contrast to Rosetta NGK, which performs very well on the positive control (1.33 Å), but poorly on the real-world conotoxin benchmark (3.5 Å). This is an expected result, as Rosetta NGK is primarily designed to complete missing regions in otherwise high-quality crystal structures. Note that as a high-resolution protocol, Rosetta NGK does a good job in generating accurate local linker conformations; it is the global positioning of the linker onto the rest of the conotoxin structure where Rosetta NGK is outperformed by PEP-Cyclizer.

PEP-Cyclizer was then applied to the identification of linkers for the design of two head-to-tail cyclized peptides, the first based on an Nrf2 fragment, the second on urotensin II. There are several significant differences between the two cases.

First, the Nrf2 peptide has been previously crystallized in linear form, in interaction with Keap1 [52]. The known linear Keap1-bound peptide structure provides a straightforward starting point for PEP-Cyclizer to design a cyclic peptide. In contrast, for UII, the absence of a full-length structure made it necessary to perform molecular modelling in order to generate an ensemble of linear peptide structures.

Second, for Nrf2, its interaction with Keap1 provides an *in vitro* test, as it is possible to measure the affinity of binding of various peptide candidates using MST. In contrast, UII binds to a membrane receptor, making an *in vitro* test extremely challenging. However, a functional assay for UII is available in the form of an intracellular calcium concentration response to extracellular UII binding.

Third, for UII, the intial single cyclic peptide structure ensemble showed considerable conformational diversity, consistent with the knowledge that although the NMR structure of the disulfide-bridged core of UII is well-defined, the flanking linear extremities are very flexible [63–66]. Depending on the experimental environment (water or membrane mimetic micelles) and temperature, distinct conformations are stabilized within the disulfide-bridged core involving different sets of intramolecular hydrogen bonds. This complicates the design of a head-to-tail cyclic peptide. Nevertheless, the fact that UII is a very potent molecule indicates that the loss of peptide conformational entropy upon binding is limited. In contrast, the Nrf2 peptide does not contain any disulfide bridges, opening up the possibility that head-to-tail cyclization may lead to rigidification, hence reduced loss of peptide conformational entropy upon binding, hence improved affinity. Together with the known linear conformation, this makes Nrf2 a relatively straightforward case; on the other hand, for UII, a cyclized peptide with demonstrated functional activity may have direct therapeutic relevance.

The differences between the two cases are reflected in the results. In both cases, using a linker predicted by our approach, a head-to-tail cyclized peptide was synthesized and its activity validated experimentally. For Nrf2, this resulted in an *in vitro* 26-fold increase of binding affinity, indicating that successful rigidification took place. In contrast, for UII, already rigidified through disulfide bond cyclization, head-to-tail cyclization resulted in a loss of activity. However, as UII is very potent to begin with (0.7 *nM*), the cyclized UII is still a very strong UT ligand (46 *nM*). Peptide cyclization still has the expected benefit of increased stability against enzymatic degradation, paying a very modest price in terms of binding affinity. Also note that in this study, we have favored short size linkers and not explored the impact of linker size.

The robustness of PEP-Cyclizer to conformational change is also apparent for the prediction of linker sequences. For UII, sequence prediction was performed on two different structure ensembles that were of different origin and highly divergent, with very similar results. Note that it is inherently complicated to evaluate linker sequence predictions, as we only have a few positive cases and no negatives, *i*.*e*., we normally do not know that a sequence does *not* cyclize. In addition, we must stress that PEP-Cyclizer proposes linker candidates based on likely sequence and structure only; In contrast, it cannot predict the effectiveness of the in vitro synthesis, nor the peptide conformational stability. Future research will focus on the prediction of the most likely length of the linker sequence, for which the current protocol does not show significant predictive power. Still, the result that experimental sequences were on average better ranked shows that PEP-Cyclizer has at least some predictive power. More importantly, the activity of the head-to-tail cyclic UII peptide LV-4130 demonstrates that PEP-Cyclizer has direct practical ability in cyclic peptide-based drug design.

Finally, although our approach shows progress for head-to-tail peptide cyclization, due to its data mining approach, it still comes with a significant limitation in terms of the nature of the types of amino acids that can be considered. The vast majority of the protein structures available in the Protein Data Bank contain only 20 standard L-amino acids. D- and unusual amino acids types that are now available for routine synthesis are currently too fewly represented to be considered by our protocol, another challenge to further consider in the future.

## Materials and Methods

In this section we explain the details of the PEP-Cyclizer protocol, that is an extension of a former data-based approach using remote or unrelated structures for loop modelling [49, 50]. Starting from the geometry of flank residues, *i*.e., four residues before and four residues after the loop of interest, it first mines a structure database and identifies all possible candidates. It then integrates a filtering step, and in the end, ranks the candidates and proposes a final set of top models (structures or sequences). PEP-Cyclizer implements two complementary and new functionalities: (*i*) guessing the linker sequence and (*ii*) modelling the conformation of the linker. The details of those functionalities are depicted in **Figure 1**, and explained in the followings.

### Structure Database

We employed two different structure databases. The first one is the database to search for linker candidates, which contains the entire set of protein structures available in the PDB. In March 2017, it consisted of 123,417 PDB entries, corresponding to 338,613 chains in total. The second database, is the one to search for linker sequences and contains the entire set of protein structures available in the PDB70. For every database, each chain was split into segments that correspond to consecutive regions separated by gaps or non-standard residues, but accepting seleno-methionines. This led to two databases with 758,143 and 172,693 protein segments, respectively.

### Test sets

To validate our approach, we have searched for cases for which both structures of the un-cyclized and cyclized peptides are available. Backbone cyclization has been applied to few conotoxins, as reported in [67], and to the best of our knowledge, the structures (NMR/Xray) of only five engineered cyclic conotoxins for which the structure of the uncyclized form exists have been deposited in the PDB database [61]. For one of the cases, two structures of the open form have been deposited in PDB (1m2c and 1mii), and their structures deviate by 1Å, and we have included both structures in our test set. For 3 additional peptides, the structure of the un-cyclized conformation and information about successful linkers are available. **Table 4** reports the details of those studied cases. Of note, the structure of all the linear and cyclic peptides in this test set have been determined using NMR, at the exception of one case (4ttl) for which it has been solved by X-ray crystallography.

Since all the structures of the un-cyclized forms of the peptides have been determined using NMR and have *N*_*uncyclized*_ conformations, we have performed the head-to-tail cyclization starting from all *N*_*uncyclized*_ NMR conformations. The final predictions for the cyclized forms of the peptides have been in turn compared with all the *N*_*cyclized*_ conformations of the cyclized structures. **Table 1** summarises the average local and global 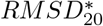 (best out of top 20) and 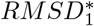 (top 1) values obtained for each linker (averaged over *N*_*uncyclized*_ conformations).

### Input preparation and candidate search

We consider head-to-tail cyclization as a loop modelling problem, where the loop flanks are the first and the last four residues in the N-terminus and C-terminus, respectively. Accordingly, the minimum acceptable size for the input linear peptide is 8 residues. We then, switch the flanks and search for linker candidates that match those flanks. We employ the method that was previously introduced to mine the database using a Binet-Cauchy (BC) kernel and a Rigidity score [68] (detail in **Supporting Materials**).

### Candidate filtering

In most cases the number of candidates returned by BCLoopSearch is too large to be tractable, which implies to limit their number. Different filters were sequentially applied in our protocol for each mode of prediction:

### Modelling the conformation of the linker

- **Sequence similarity:** The sequence similarity of a linker candidate with the query linker sequence using BLOSUM62 score. Candidates with negative scores were discarded.
- **Geometrical clustering:** We used the python Numpy library to measure the pairwise distances (RMSD) between all the candidates [69]. In addition, we used the python Scipy package to perform hierarchical clustering [70]. A RMSD cutoff of 1Å was used to group similar linker candidates. To consider memory constraints, we applied an iterative clustering over subsets of 25,000 candidates, until at most 25,000 clusters were obtained.
- Finally, one representative linker candidate with the highest sequence similarity to the query linker was selected for each cluster. The computational time of our clustering protocol is in the range of 1-5 minutes, however it depends directly on the number of candidates detected by BCLoopSearch. In extreme cases, the needed time may increase up to 10-15 minutes.
- **Local conformation:** Previously, Shen et al. have shown that local conformation profiles predicted from sequence and profile-profile comparison can be employed to accurately distinguish similar structural fragments [71]. Consequently, we pre-computed a collection of profiles for all the protein chains in the structure dataset, and for all proteins of the test sets. For each linker candidate, it is thus possible to extract the sub-profiles *P* and *Q*, corresponding to the query and candidate linker, and to measure the Jensen Shannon divergence (*JS*(*P, Q*)) between these profiles:

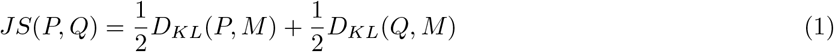

where *M* corresponds to 1*/*2(*P* + *Q*) and *D*_*KL*_ is the Kullback-Leibler divergence:

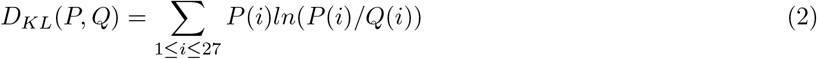

*P* (*i*) is the probability of SA letter *i*. Then we measured the average Jensen Shannon divergence (*JSD*) over the paired series of query and candidate profiles:

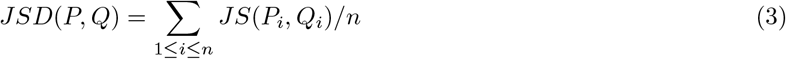

where *P*_*i*_ and *Q*_*j*_ are the two profiles corresponding to positions 1 to L on the query and candidate linker sequences. Note that a *JSD* of 0 indicates a perfect identity of the profiles. This procedure was applied on each linker candidate and those with a *JSD >* 0.40 were discarded from the remaining set.

- **steric clash detection:** After modelling the complete structure, models with steric clashes were discarded considering the C_*α*_ distance between linker residues and other residues of the protein, using a cutoff value of 3 Å.

### Predicting the linker sequence

- **Sequence similarity:** If sequence constraints are given, a subset of sequences that represent at least 50% sequence identity to any of the constraint amino acid types, regardless of their position, are kept.
- **Local conformation:** Measuring the local conformation of flanks (query and candidate flanks) and discarding candidates with flank *JSD >* 0.40.

### Sequence constraints

Throughout the study, linker sequences were predicted using the following sequence constraints. At each position of the linker, the set of amino acids of the entire experimental linker was considered - for instance, for the RGD linker of 2ew4, the amino acids Arg, Gly and Asp were considered at all three positions, *i*.*e*., 3^3^ different linker sequences are possible.

### Model building

Final energy minimisation was conducted using Gromacs 2018 [72], the CHARMM36m force field [73] and the steepest descent algorithm for 1000 steps. All bonds were constrained using the LINCS algorithm. The particle mesh Ewald algorithm was used to handle electrostatics with a 10 Å cutoff for the short-range part and a grid spacing of 1.2 Å for the long-range contribution in reciprocal space. The Verlet buffer scheme was used for non-bonded interactions, the neighbour list was updated every 20 steps.

### Model selection

To rank the models, we considered the RMSD of the flanks. In case of conformation modelling, our procedure returns a maximum of 20 models with the lowest *flank RMSD* score. And for sequence guessing, it returns a set of 30 sequences with the lowest *flank RMSD* score. From this set and considering the sequence constraints, we apply the sequence inference procedure (as explained below) to propose final set of likely sequences for the linker.

### Candidate sequence inference

To draw candidate sequences given the sequences of the candidate linkers identified, we have used a forward-backtrack procedure. One advantage of such a procedure is to provide both sequences and their likelihood. The probabilities 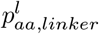 of observing each amino acid type *aa* at position *l* of the *linker* can be estimated from the amino acid sequences of the candidate linkers satisfying the condition of peptide cyclization. However, when a reduced number of amino acids is considered at a given position, these estimates can be performed on a rather low number of sequences. Consequently, we have estimated pseudo-frequencies, with 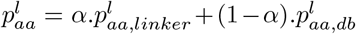 where *α* is a value between 0 and 1, and 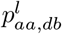 is the frequency of amino acid type *aa* as observed in a large collection of sequences named *db*. For *db*, we have considered the sequences of the loops of 123,417 PDB entries (758,143 protein segments), identified using the procedure described in [49]. Alternatively, we have also considered *db*_*s*_, which corresponds to the subset of *db* corresponding to a loop size of *s*. Transition probabilities have been estimated similarly. Pseudo transition probabilities *p*(*aa*^*l*^*/aa*^*l−*1^) were estimated as 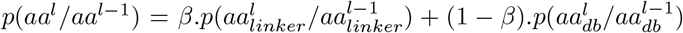 where *β* is a value between 0 and 1. Given estimates of 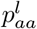 and *p*(*aa*^*l*^*/aa*^*l−*1^) we have used the forward-backtrack algorithm to infer series of amino acids that fit best the estimates. We prefer such procedure to for instance the *viterbi*_*kbest*_ procedure that, in our experience [74], usually returns less diverse sequences.

### Linker quality assessment

To assess the quality of the final linker structures, we use the global RMSD of the linker candidates main chain heavy atoms (N, C, C*α* and O), *i*.*e*., the modeled cyclic peptides are superimposed on the native structure excluding the linker region, then the RMSD is calculated over the linker.

### Statistical testing

To test the prediction of linker sequences of the conotoxin benchmark, the rank of the experimental linker sequences were determined. To avoid pseudo-replication, five duplicate cyclic sequences were eliminated; using the remainder of the benchmark, the overall ranking of the experimental linker sequences was tested for statistical significance. With the total number of linker sequences varying from case to case, and many instances of tied ranks, it was not feasible to compute an analytical p-value based on hypergeometric distributions. Instead, random ranks were simulated by sampling from flat rank distributions, converted to percentiles, and it was evaluated how often the overall mean percentile was better than the observed mean percentile (37.4) for the experimental linker sequences. This was the case in 2518/100000 random simulations, *i*.*e*., a p-value of 0.025.

### Comparison with other approaches

In this work we compare the performance of our linker modelling protocol with the Rosetta NGK [51]. The Rosetta NGK runs were performed using the protocol provided by [51], and Rosetta energy values were employed for ranking the models. Considering the fact that Rosetta NGK is not designed for peptide cyclization, we converted the head-to-tail cyclization to loop modelling, by breaking every peptide into two segments and switching the two.

### Experimental tests

To test our procedure, we have searched for linear peptides known to interact with a protein, and for which some quantification of the impact of the cyclization can be done. Since the aim of our approach to design linkers is to perturb as few a possible the peptide conformation, while possibly rigidifying it, a straightforward possibility was to search the Protein Data Bank to identify peptide protein complexes solved at high resolution, for which the peptide does not contain unusual amino acids, and for which the protein is available commercially so that it is possible to undergo affinity measurements between the different peptides and the protein. Two such systems were identified and correspond respectively to PDB entries 1×2r and 2qos. The first one corresponds to the interaction of a fragment of Nrf2 of 9 amino acids (sequence: LDEETGEFL) in interaction with Keap1, an interaction spotted in the context of lung cancer. The second one corresponds to the C8 binding site, a peptide of 11 amino acids (sequence: LRYDSTAERLY) interacting with complement protein C8. However, this peptide led to aggregation when tested for MST and did not lead to any exploitable results. A final system corresponds to urotensin II, a peptide of 11 amino acids (sequence: ETPDCFWKYCV) known to interact with a membrane receptor, for which functional tests of ligand-stimulated intracellular calcium response are available commercially.

### Urotensin II model generation

Two sets of 3D models were used. The first one was generated using PEP-FOLD server [35], a *de novo* approach to peptide structure prediction. Five independent runs of 3D generation (100 models) were run, and five models showing closed disufide bonds in the PEP-FOLD coarse grained representation were then submitted to refinement using MD, with the aim to stabilize the disulfide bond in the all atom representation. The model topology was created using the Gromacs pdb2gmx command, which did not include the disulfide bond. The topology was further modified to include the disulfide bond parameter using the gromacs py library [75]. Simulations were performed using the CHARMM-36 force field [76] and the TIP3P model for water. The Gromacs 2018 software [72] was used to run the simulations. The five models were minimized two times for 10,000 steps with the steepest descent algorithm. During the first minimisation the bonds were not constraints, as in the second and following steps, all bonds were constrained using the LINCS algorithm. The five models were solvated in a water box and roughly 150 *mM* of NaCl. Systems were again minimized in two similar steps, then equilibrated in three successive steps, *(i)* 100 *ps* with position restraints of 1000 *kJmol*^*−*1^*nm*^*−*2^ applied on the peptide heavy atoms and an integration time step of 1 fs *(ii)* 500 *ps* with position restraints of 1000 *kJmol*^*−*1^*nm*^*−*2^ applied on the peptide C*α* atoms, the integration time step was fixed to 2 *fs (iii)* 1 *ns* with position restraints of 100 *kJmol*^*−*1^*nm*^*−*2^ applied on the C*α* atoms. Production runs were finally computed for 100 *ns*. The five 100 *ns* trajectories were then analysed using MDAnalysis library [77]. PCA of backbone atoms coordinates were computed and the fifteen first components were used to cluster the coordinates. The clustering DBSCAN algorithm [78] was used using a min sample of 20, and sigma value of 5. A total of 13 clusters was identified, the cluster centroids were chosen by taking the closest element in terms of RMSD to the average cluster structure. The conformations generated using this protocol are available as supplementary information. All models underwent sequence guessing to cyclize the peptide.

Another set of models was kindly provided by D. Chatenet and co-workers, at INRS Quebec, Canada. It consists of a set of 8 representative structures of UII displaying the heterogeneous conformational ensemble of this peptide. The three-dimensional structure of UII was generated from the sequence using the pdbutilities server https://spin.niddk.nih.gov/bax/nmrserver/pdbutil/. System preparation and MD simulations were performed using AMBER v16 [79] and the ff14SB force field [80]. Simulations were performed at 300 K under constant energy (NVE) conditions using a 2 *fs* timestep. The peptide was solvated using the SPC(E) water model in a rectangular box with periodic boundary conditions. The system was neutralized through the addition of counter ions (*Na*^+^). The pre-processing steps were followed by equilibration steps, as described previously [81]. All simulations were performed using the GPU-enabled version of the AMBER simulation engine pmemd. A Particle Mesh Ewald cutoff of 8 Å was used for the GPU-enabled simulations [82]. The peptide was simulated for a total of 100 *ns*. Representative structures were selected by clustering simulation ensembles obtained from the MD simulation trajectory. Clustering was performed using the hierarchical agglomerative approach with an epsilon cutoff of 3 Å, which represents the minimum distance between the clusters.

### Peptide synthesis and functional test

Linear peptide precursors of human UII (LV-4130, LV-4131; LV-4132 and LV-4133) and human/mouse *Nrf* 2_(76*−*85)_ (LV-5562 and LV-5565) cyclic analogues were synthesized by Fmoc solid phase methodology on a Liberty microwave assisted automated peptide synthesizer (CEM, Saclay, France) using the standard manufacturer’s procedure at 0.1 mmol scale on a preloaded Fmoc-Asp(Wang resin)-ODmab as previously described [83]. Reactive side chains were protected as follow: Thr, Tyr, tert-butyl (tBu) ether; Glu, tert-butyl (OtBu) ester; Lys, Trp, tert-butyloxycarbonyl (Boc) carbamate; Cys, trityl (Trt) thioether. After completion of the chain assembly, the C-terminal Dmab protective group was selectively removed by addition of a solution of 2% hydrazine in DMF for 3 min [84]. Treatment was repeated twice and the resin was washed with DMF and DCM. Head-to-tail cyclization was performed on-resin by in situ activation of the free carboxyl group with BOP (1.1 eq) and DIEA (10 eq) in 10 mL of DMF at room temperature for about 72 h. The head-to-tail cyclic peptides were deprotected and cleaved from the resin by adding 10 mL of the mixture TFA/TIS/H2O (9.5:0.25:0.25) for 180 min at room temperature. After filtration, crude peptides were washed thrice by precipitation in TBME followed by centrifugation (4500 rpm, 15 min). The synthetic peptides were purified by reversed-phase HPLC on a 21.2 × 250 mm Jupiter C18 (5 *µm*, 300 Å) column (Phenomenex, Le Pecq, France) using a linear gradient (10-50% over 45 min) of acetonitrile/TFA (99.9:0.1) at a flow rate of 10 mL/min. The disulfide bridge of UII analogues was then formed by treatment of the head-to-tail cyclic peptides with a mixture of N-chlorosuccinimide (1.05 eq) in 20 mL of *H*_2_*O/CH*_3_*CN* (1:1) for 30 min at room temperature as previously described [85]. The resulting bicyclic UII analogues were purified as described above with a 20-60% linear gradient. The purified peptides were finally characterized by MALDI-TOF mass spectrometry on a ultrafleXtreme (Bruker, Strasbourg, France) in the reflector mode using *α*-cyano-4-hydroxycinnamic acid as a matrix. Analytical RP-HPLC, performed on a 4.6 × 250 mm Jupiter C18 (5 *µm*, 300 Å) column, indicated that the purity of all peptides was *>* 95%. Purity analyses of 4 peptides are reported in **supplementary Figure S1**.

### Intracellular calcium assay

Ligand-stimulated intracellular calcium responses were measured at the human UT receptor expressed in transfected CHO cells using a fluorimetric detection method according to Eurofins-Cerep (catalog ref. G099-1376) and EuroscreenFast (catalog ref. FAST-0540A) standard assay protocols. The assays were performed in duplicate or triplicate. The results were expressed as a percent of human UII response at its *EC*_100_ concentration and plotted using Prism software (GraphPad, San Diego, CA).

### Microscale thermophoresis

Keap1 (His-tagged) was purchased from Tebu-Bio (Le Perray en Yvelines, France). This His-tagged protein was labeled using the Labeling Kit RED-tris-NTA 2nd Generation Monolith (MO-L018, NanoTemper Technologies GmbH, Germany). According to the manufacturer’s instructions we have first evaluated the affinity of the dye toward the His-Keap1, by preparing 16 dilution points of the his-Keap1 protein in PBS-T (consisting of PBS supplied with the labelling kit to which has been added 0.05% of tween 20) (from 4 *µM* to 0.12 *nM* ; 10 *µl* each) and finally adding 10 *µl* of the RED-tris-NTA 2nd generation dye (50 *nM*) (in PBS-T) in each tube. After 30 min of incubation at room temperature, the affinity (Kd) of the RED-tris-NTA 2nd generation dye for Keap1 His-tag was measured using the Monolith NT.115Pico instrument. As recommended by the manufacturer, since we measured a Kd at 18 *nM*, we have chosen to label His-Keap1 as follow for all experiments dedicated to the affinity measurement of His-Keap1 with the different putative ligand. Briefly, we adjusted 90 *µl* of the Keap1 concentration at 362 *nM* and we added 90 *µl* of RED-tris-NTA 2nd generation dye (100 *nM*). After 30 min of incubation at room temperature, the sample was centrifugated (10 min; 4 ^*◦*^C; 15,000 g) and then the supernatant was collected for binding assay. Concerning the binding assay experiments, the linear *Nrf* 2_(76*−*85)_ peptide, LV-5562 and LV-5565 were diluted in pure *H*_2_*O* at 600 *µM*. 20 *µl* of these stock solutions were placed in tube 1 and a series of 1:1 dilution were prepared in PBS-T, in order to obtain a ligand concentration ranged from 600 µM to 18.32 *nM* (16 points). Afterward, 10 *µl* of the labelled protein (100 *nM*) was added in each 16 tubes containing the ligand at the 16 different concentration. Finally, for MST measurement the final concentration was 300 *µM* to 9.16 *nM* of ligand and 50 *nM* of His-Keap1 labelled protein. After this preparation, each solution was filled into Monolith NT Standard Capillaries. After loading the 16 capillaries into the Monolith NT.115Pico instrument, a scan of the fluorescence count for all capillaries was carried to check for consistent fluorescence while confirming the absence of ligand induced fluorescence changes or adsorption. MST was measured using a Monolith NT.115Pico instrument at an ambient temperature of 25 ^*◦*^C with 5 s/30 s/5 s laser off/on/off times, respectively. Instrument parameters were adjusted with 5% LED power and 40% MST power. Data of at least three independently pipetted measurements were analyzed (MO.Affinity Analysis software version 1.5.41, NanoTemper Technologies GmbH) using FHOT at 5 sec. The data were fitted using the law of mass action from GraphPad Prism version 5, and MST figures were generated using MO.Affinity Analysis.

## Acknowledgements

ANR-10-BINF-0003 (BipBip); ANR-14-2011-IFB; INSERM [U 1133]; Ressource Parisienne en Bioinformatique Structurale (RPBS). The authors thank David Chatenet, Nicolas Doucet and Chitra Narayanan, INRS, Quebec, Canada for communicating sample conformations of UII.

## Table of Contents graphic

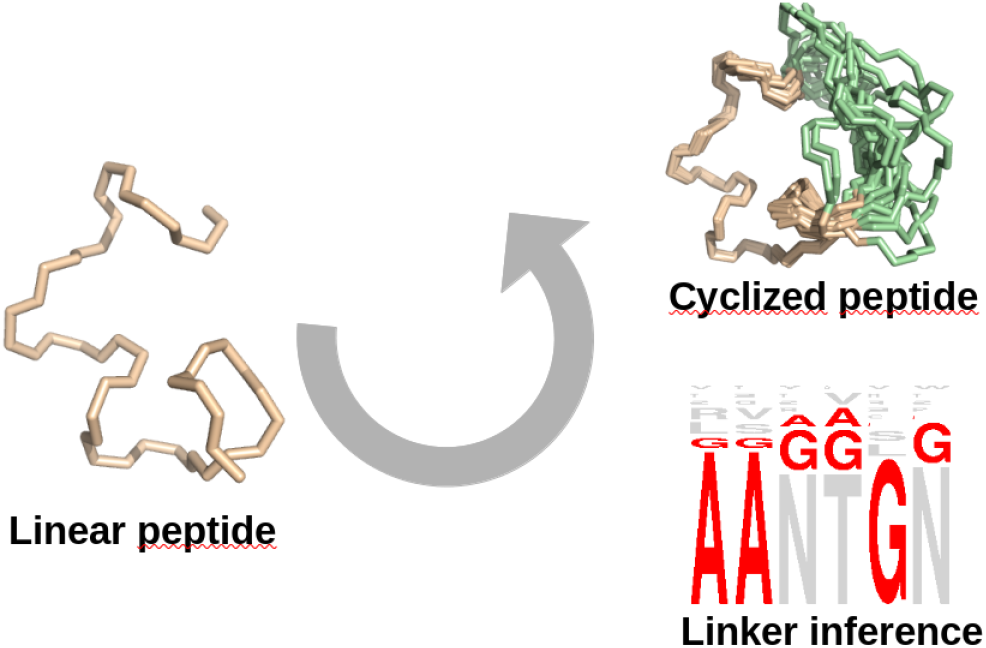

