## Supplementary information for "Head-to-tail peptide cyclization: new directions and application to urotensin II and Nrf2"

### A novel computational method for head-to-tail peptide cyclization: application to urotensin II and Nrf2 Supplementary Materials

Yasaman Karami[1], Samuel Murail[1], Julien Giribaldi[2], Benjamin Lefranc[3], Florian Defontaine[4], Olivier Lesouhaitier[4], Jérôme Leprince[3], Sjoerd J. de Vries[1,*], Pierre Tufféry[1,*]

[1] Université Paris Cité, CNRS UMR 8251, INSERM U1133, Paris, France.
[2] Institut des Biomolécules Max Mousseron, UMR 5247, Université de Montpellier-CNRS, Montpellier, France.
[3] Université de Rouen Normandie, INSERM U1239 NorDiC, Neuroendocrine, Endocrine and Germinal Differentiation and Communication, INSERM US51 HeRacLeS, Rouen, France.
[4] Université de Rouen Normandie, UR CBSA, Research Unit Bacterial Communication and Anti-infectious Strategies, Evreux, France.

#### Database search

We previously introduced the BCLoopSearch protocol, to mine large protein structure datasets and retrieve loop candidates, given two disjoint fragments (loop flanks) [1]. It is based on a Binet-Cauchy (BC) kernel and a Rigidity score:

$$BC(X,Y) = \frac{det(X^TY)}{\sqrt{det(X^TX)det(Y^TY)}} \tag{1}$$

where $X$ and $Y$ are $C_\alpha$ coordinates of the flanks and dataset fragments, respectively and they are centered at the origin. Note that a BC score of 1 indicates a perfect match. *Rigidity* score $R(X,Y)$ is defined as:

$$R'(X,Y) = max_{1\leq i\leq N}|||X_i - Y_i||| \tag{2}$$

$$R(X,Y) = max\{R'(X,Y), |||X_N - X_1\| - \|Y_N - Y_1\||\} \tag{3}$$

where $X_i$ and $Y_i$ are $C_\alpha$ coordinates of the $i$th residues of the flanks and dataset fragments and $||.||$ is the euclidean norm. Rigidity score is the maximum variation of intra-distances between: *(i)* residues and geometric center and *(ii)* intra-distances between terminal $C_\alpha$. In addition, we also measured the RMSD between query and candidate flanks for the fragments returned. In total, four cutoffs values related to *(i)* flank size, *(ii)* flank BC score, *(iii)* flank Rigidity and *(iv)* flank RMSD, have been considered to limit the number of loop candidates. In this study we used: a flank size of 4 residues, Rigidity $\leq$ 2.5, flank RMSD $\leq$ 4 Å and the minimal flank BC score cutoff of 0.8.

### CyBase benchmark

CyBase (http://www.cybase.org.au/) [2, 3] provides a set of existing naturally occurring cyclic peptides. Presently, 64 3D structures of cyclic peptides from 25 different species are reported. We applied a filtering step on the list to keep only those that are *i)* head-to-tail cyclized, *ii)* without modified amino acids and *iii)* not identical (filtering out entries with identical sequences), resulting in a final set of 35 structures. Residues from the N- and/or C-terminal extremities of each cyclic peptide were removed to generate linear peptides (here by N- and C-terminal extremity, we refer to the head and tail residues from the sequence). We considered all possible combinations of truncating two to seven residues from the N- and/or C-termini (*i.e.*, removing two residues from N-terminus or two residues from C-terminus or one residue from each side), generating 33 different linear peptides from every cyclic target. We also excluded the cases where the size of generated linear peptide was less than 8 residues, that is size limit of our protocol. Finally we obtained a total of 1147 linear peptides, where the corresponding linkers are in the range of 2-7 residue long. The details of those structures are reported in **Supplementary Table S1**.

Table S1: **The list of cyclic structures from CyBase.** Structures with identical sequences were discarded and only one representative was considered. For each cyclic peptide, we generated a total of 33 linear peptides by truncating two to seven residues from N- and/or C-term. The total number of linear peptides for each target, as well as those modelled with PEP-Cyclizer and Rosetta NGK are reported.

| Protein name | Class | Type | PDBcode | size | #linkers | #linkers (PEP-Cyclizer) | #linkers (NGK) |
|---|---|---|---|---|---|---|---|
| kalata-B1 | Cyclotide | NMR | 1NB1 | 29 | 33 | 33 | 33 |
| kalata-B1 | Cyclotide | NMR | 1K48 | 29 | 33 | 33 | 32 |
| kalata-B1 | Cyclotide | NMR | 1KAL | 29 | 33 | 33 | 32 |
| [P20D,V21K]-kalata-B1 | Cyclotide | NMR | 2F2I | 29 | 33 | 33 | 33 |
| [W19K,-P20N,-V21K]-kalata-B1 | Cyclotide | NMR | 2F2J | 29 | 33 | 33 | 33 |
| kalata-B2 | Cyclotide | NMR | 1PT4 | 29 | 33 | 33 | 33 |
| kalata-B5 | Cyclotide | NMR | 2KUX | 30 | 33 | 32 | 33 |
| kalata-B7 | Cyclotide | NMR | 2JWM | 29 | 33 | 33 | 33 |
| kalata-B7 | Cyclotide | NMR | 2M9O | 29 | 33 | 33 | 33 |
| kalata-B8 | Cyclotide | NMR | 2B38 | 31 | 33 | 33 | 33 |
| kalata-B12 | Cyclotide | NMR | 2KVX | 28 | 33 | 32 | 33 |
| cycloviolacin-O1 | Cyclotide | NMR | 1NBJ | 30 | 33 | 32 | 33 |
| cycloviolacin-O1 | Cyclotide | NMR | 1DF6 | 30 | 33 | 33 | 33 |
| cycloviolacin-O2 | Cyclotide | NMR | 2KNM | 30 | 33 | 32 | 33 |
| cycloviolacin-O14 | Cyclotide | NMR | 2GJ0 | 31 | 33 | 33 | 33 |
| MCoTI-II | Squash-trypsin-inhibitor | XRAY | 4GUX | 34 | 33 | 33 | 33 |
| circulin-A | Cyclotide | NMR | 1BH4 | 30 | 33 | 32 | 33 |
| circulin-B | Cyclotide | NMR | 2ERI | 31 | 33 | 33 | 33 |
| kB1[GHFRWG;23-28] | Cyclotide | NMR | 2LUR | 29 | 33 | 32 | 32 |
| [Ala1,15]kB1 | Cyclotide | NMR | 1N1U | 29 | 33 | 33 | 33 |
| des(24-28)kB1 | Cyclotide | NMR | 1ORX | 24 | 33 | 33 | 32 |
| SFTI-1 | BBI-like-trypsin-inhibitor | XRAY | 3P8F | 14 | 25 | 25 | 25 |
| Ent-AS-48 | Bacterial | XRAY | 1O82 | 70 | 33 | 33 | 33 |
| vhl-1 | Cyclotide | NMR | 1ZA8 | 31 | 33 | 33 | 33 |
| vhl-2 | Cyclotide | NMR | 2KUK | 30 | 33 | 33 | 33 |
| varv-peptide-F | Cyclotide | NMR | 2K7G | 29 | 33 | 33 | 33 |
| varv-peptide-F | Cyclotide | XRAY | 3E4H | 29 | 33 | 33 | 33 |
| BiKK | BBI-like-trypsin-inhibitor | NMR | 2BEY | 16 | 33 | 33 | 33 |
| RTD-1 | Primate | NMR | 1HVZ | 18 | 33 | 33 | 33 |
| palicourein | Cyclotide | NMR | 1R1F | 37 | 33 | 33 | 33 |
| vhr1 | Cyclotide | NMR | 1VB8 | 30 | 33 | 31 | 33 |
| tricyclon-A | Cyclotide | NMR | 1YP8 | 33 | 33 | 33 | 33 |
| Cter-M | Cyclotide | NMR | 2LAM | 29 | 33 | 33 | 33 |
| MCo-PMI | Squash-trypsin-inhibitor | NMR | 2M86 | 51 | 33 | 33 | 31 |
| Carnocyclin-A | Bacterial | NMR | 2KJF | 60 | 33 | 33 | 33 |
| total | | | | | 1147 | 1141 | 1141 |

We applied both our protocol and Rosetta NGK to the CyBase test set to model all the linkers. Over the 1147 cases, both our data-mining and Rosetta NGK failed to model the linker for 6 different cases (0.5%). In fact, our protocol identified candidates in all cases, but discarded all the candidates with a correct geometry but a non satisfactory sequence similarity in 6 cases. Thus overall, in terms of ability to identify linkers, the data-mining strategy seems to perform as well as a pure *ab initio* procedure. Then, we compared both protocols using the 1135 over 1147 (99%) cases for which the linker could be modelled by both methods. All heavy backbone atoms (N, C, C$\alpha$, O) were considered. The local RMSD corresponds to RMSD obtained by superposing the model linker on the native conformation using a best fit procedure, whereas the global RMSD corresponds to RMSD observed after superposing the linear part of the peptide (*i.e.*, without the linker). The best RMSD over the top 20 predictions by each method were retained.

Table S2: **The RMSD values for all the linkers of each structure from CyBase.** The average local and global RMSD values are measured over the backbone atoms (N, C, C$\alpha$, O) for the linkers modelled by both PEP-Cyclizer and Rosetta NGK. For each cyclic peptide, we generated a total of 33 linear peptides by truncating two to seven residues from N- and/or C-terminal extremities. For each target, the number of linear peptides that were cyclized by both PEP-Cyclizer and Rosetta NGK are reported (out of the total 33 linkers).

| **Protein name** | **number of linkers** | local RMSD (Å) | | global RMSD (Å) | |
|---|---|---|---|---|---|
| | | **PEP-Cyclizer** | **NGK** | **PEP-Cyclizer** | **NGK** |
| kalata-B1 | 33 | 0.53±0.22 | 0.56±0.37 | 1.17±0.45 | 1.05±0.46 |
| kalata-B1 | 32 | 0.69±0.34 | 0.70±0.57 | 2.05±1.40 | 1.77±2.43 |
| kalata-B1 | 32 | 0.71±0.26 | 1.04±0.55 | 1.45±0.47 | 1.71±0.96 |
| [P20D,V21K]-kalata-B1 | 33 | 0.65±0.37 | 0.52±0.38 | 1.40±0.70 | 0.99±0.77 |
| [W19K,-P20N,-V21K]-kalata-B1 | 33 | 0.54±0.28 | 0.71±0.40 | 1.31±0.70 | 1.24±0.65 |
| kalata-B2 | 33 | 0.49±0.17 | 0.40±0.38 | 1.12±0.47 | 0.73±0.66 |
| kalata-B5 | 32 | 0.74±0.45 | 0.45±0.57 | 1.83±1.53 | 0.88±1.54 |
| kalata-B7 | 33 | 0.58±0.19 | 0.82±0.65 | 1.15±0.32 | 1.13±0.73 |
| kalata-B7 | 33 | 0.79±0.36 | 0.56±0.58 | 2.11±1.52 | 1.45±2.06 |
| kalata-B8 | 33 | 1.12±0.57 | 1.14±0.61 | 2.57±1.13 | 2.11±1.22 |
| kalata-B12 | 32 | 0.74±0.38 | 0.69±0.37 | 1.57±1.07 | 1.40±1.32 |
| cycloviolacin-O1 | 32 | 1.08±0.68 | 0.43±0.17 | 2.83±1.98 | 0.91±0.33 |
| cycloviolacin-O1 | 33 | 0.98±0.43 | 1.15±0.43 | 2.14±1.24 | 1.99±1.17 |
| cycloviolacin-O2 | 32 | 0.75±0.43 | 0.38±0.39 | 2.07±1.76 | 0.75±1.30 |
| cycloviolacin-O14 | 33 | 0.94±0.57 | 0.79±0.89 | 2.61±1.91 | 2.01±2.54 |
| MCoTI-II | 33 | 0.73±0.43 | 0.31±0.44 | 1.70±0.78 | 0.64±1.13 |
| circulin-A | 33 | 0.97±0.34 | 0.88±0.34 | 2.37±0.96 | 1.59±0.87 |
| circulin-B | 33 | 1.02±0.44 | 0.37±0.17 | 2.22±0.91 | 0.61±0.30 |
| kB1[GHFRWG;23-28] | 32 | 1.08±0.45 | 1.16±0.40 | 2.61±1.23 | 2.18±0.87 |
| [Ala1,15]kB1 | 33 | 0.93±0.39 | 0.88±0.33 | 1.94±0.73 | 1.22±0.51 |
| des(24-28)kB1 | 32 | 1.34±0.55 | 1.60±0.64 | 2.44±0.81 | 2.88±1.37 |
| SFTI-1 | 25 | 0.50±0.41 | 0.22±0.14 | 1.48±1.19 | 0.57±0.48 |
| Ent-AS-48 | 33 | 0.54±0.29 | 0.14±0.08 | 1.16±0.53 | 0.23±0.10 |
| vhl-1 | 33 | 0.87±0.46 | 0.36±0.27 | 1.88±1.01 | 0.66±0.32 |
| vhl-2 | 33 | 0.53±0.25 | 0.71±0.74 | 1.24±0.52 | 1.11±0.95 |
| varv-peptide-F | 33 | 0.61±0.42 | 0.51±0.64 | 1.86±1.62 | 1.24±2.12 |
| varv-peptide-F | 33 | 0.50±0.21 | 0.41±0.28 | 1.17±0.52 | 0.79±0.35 |
| BiKK | 33 | 0.64±0.47 | 1.05±0.75 | 1.64±1.27 | 1.87±1.36 |
| RTD-1 | 33 | 0.72±0.37 | 0.74±0.43 | 1.98±0.84 | 1.77±0.94 |
| palicourein | 33 | 1.12±0.36 | 1.11±0.32 | 2.24±1.02 | 1.93±0.59 |
| vhr1 | 31 | 0.97±0.58 | 0.61±0.34 | 2.36±1.59 | 1.23±0.48 |
| tricyclon-A | 33 | 0.79±0.30 | 0.68±0.48 | 1.64±0.57 | 1.13±0.86 |
| Cter-M | 33 | 0.60±0.42 | 0.54±0.69 | 1.62±1.43 | 1.45±2.08 |
| MCo-PMI | 31 | 1.11±0.48 | 1.09±0.51 | 2.33±1.22 | 1.93±1.19 |
| Carnocyclin-A | 33 | 0.52±0.26 | 0.23±0.13 | 1.21±0.47 | 0.48±0.22 |

| linker size (# gaps) | | 2 (101) | 3 (139) | 4 (175) | 5(208) | 6 (241) | 7 (271) |
|---|---|---|---|---|---|---|---|
| $lRMSD_{20}$ | PEP-Cyclizer | 0.32±0.19 | 0.51±0.22 | 0.66±0.28 | 0.77±0.35 | 0.92±0.44 | 1.11±0.49 |
| | Rosetta NGK | 0.26±0.33 | 0.38±0.36 | 0.52±0.43 | 0.66±0.48 | 0.83±0.54 | 1.04±0.66 |
| $gRMSD_{20}$ | PEP-Cyclizer | 1.38±0.57 | 1.45±0.50 | 1.52±0.59 | 1.64±0.78 | 2.02±1.25 | 2.55±1.70 |
| | Rosetta NGK | 0.73±0.78 | 0.84±0.70 | 0.97±0.68 | 1.15±0.81 | 1.60±1.51 | 1.93±1.74 |
| $lRMSD_1$ | PEP-Cyclizer | 0.64±0.29 | 0.94±0.39 | 1.14±0.50 | 1.21±0.57 | 1.43±0.67 | 1.79±0.80 |
| | Rosetta NGK | 0.34±0.35 | 0.55±0.53 | 0.73±0.61 | 0.89±0.63 | 1.12±0.72 | 1.35±0.80 |
| $gRMSD_1$ | PEP-Cyclizer | 2.67±1.40 | 2.80±1.47 | 3.06±1.85 | 3.06±2.18 | 3.99±2.67 | 4.90±3.14 |
| | Rosetta NGK | 0.89±0.89 | 1.14±0.99 | 1.44±1.47 | 1.68±1.74 | 2.18±1.87 | 2.65±2.25 |

Table S3: **RMSD and ranks over the CyBase test set.** For every case the best RMSD out of top 20 and the top 1 were considered. The average and standard deviations of best local ($lRMSD_{20}$, $lRMSD_1$) and global ($gRMSD_{20}$, $gRMSD_1$) RMSD values are reported for every gap size.

Table S4: **Summary of cyclic linkers for conotoxins.** Data is collected from [4] and additional details are added from the mentioned references. The last column reports the pdb code of the available engineered cyclic peptides. linkers sequences in bold correspond to the functional variants that were considered in this study.

| name | linear peptide | linker | activity | structure | stability | pdb code |
|---|---|---|---|---|---|---|
| | | GGAAG (cMII-5) [5] | not active | not similar | - | - |
| α-Conotoxin MII | 1m2c (1mii) | **GGAAGG (cMII-6)** [5] | similar | similar | improved | **2ajw** |
| | | **GAGGAAG (cMII-7)** [5] | smiliar | similar | improved | **2ak0** |
| α-Conotoxin ImI | | A [6] | - | - | slightly improved | - |
| | | βA [6] | - | - | improved | - |
| | 1cnl | AG [6] | - | - | slightly improved | - |
| | | AGG [6] | - | - | slightly improved | - |
| α-Conotoxin Vc1.1 | 2h8s | GGAAG [7] | substantial loss | similar | - | - |
| | | **GGAAGG** [7] | similar/higher | similar | improved | **4ttl** |
| | | GAA [8] | reduced | not similar | - | - |
| | | GAAG [8] | reduced | not similar | - | - |
| α-Conotoxin RgIA | 2JUT | GAAGG [8] | reduced | similar | - | - |
| | | **GGAAGG** [8] | similar | similar | improved | - |
| | | **GGAAGAG** [8] | similar | similar | improved | - |
| | | A [9] | reduced | - | - | - |
| | | **AG** [9] | reduced | - | improved | - |
| | | AGG [9] | reduced | - | improved | - |
| | | **AGGG** [9] | reduced | - | improved | - |
| | | GGAAG [9] | reduced | - | improved | - |
| α-Conotoxin AuIB | 1mxn | GAGAAG [9] | reduced | - | improved | - |
| | 1mxp | GGAGGAG [9] | reduced | - | improved | - |
| | | **GGAA** [10] | reduced | similar | improved | - |
| | | AGAGA [10] | reduced | similar | improved | - |
| | | GGAAGG [10] | reduced | similar | improved | - |
| | | GGAAAGG [10] | reduced | - | improved | - |
| χ-Conotoxin MrIA | 2ew4 | **AG** [11] | similar | similar | improved | 2j15 |
| | | **RGD** [12] | similar | similar | improved | - |
| ω-Conotoxin MVIIA | 1mvi | **GGPG** [13] | - | - | - | - |
| Conotoxin gm9a | 1ixt | **GLP** [14] | - | similar | similar | 2mso |
| Conotoxin bru9a | - | **GLP** [14] | - | - | similar | 2msq |

Table S5: **Average ranks of the cyclic linkers for conotoxins, using forward-backtrack algorithm.**

| name | linear peptide | linker | ranks |
|---|---|---|---|
| $\alpha$-Conotoxin MII | 1m2c (1mii) | GGAAG (cMII-5) | 19/32 |
| | | **GGAAGG (cMII-6)** | 40/64 |
| | | **GAGGAAG (cMII-7)** | 8/128 |
| $\alpha$-Conotoxin ImI | 1cnl | A | - |
| | | $\beta$A | - |
| | | AG | 2/4 |
| | | AGG | 3/8 |
| $\alpha$-Conotoxin Vc1.1 | 2h8s | GGAAG | 23/32 |
| | | **GGAAGG** | 33/64 |
| $\alpha$-Conotoxin RgIA | 2JUT | GAA | 8/8 |
| | | GAAG | 14/16 |
| | | GAAGG | 28/32 |
| | | **GGAAGG** | 24/64 |
| | | **GGAAGAG** | 21/128 |
| $\alpha$-Conotoxin AuIB | 1mxn<br>1mxp | A | - |
| | | **AG** | 1/4 |
| | | AGG | 2/8 |
| | | **AGGG** | 6/16 |
| | | GGAAG | 25/32 |
| | | GAGAAG | 24/64 |
| | | GGAGGAG | 8/128 |
| | | **GGAA** | 14/16 |
| | | AGAGA | 3/32 |
| | | GGAAGG | 37/64 |
| | | GGAAAGG | 35/128 |
| $\chi$-Conotoxin MrIA | 2ew4 | **AG** | 2/4 |
| | | **RGD** | 7/27 |
| $\omega$-Conotoxin MVIIA | 1mvi | **GGPG** | 3/16 |
| Conotoxin gm9a | 1ixt | **GLP** | 8/27 |
| Conotoxin bru9a | - | **GLP** | - |

Table S6: **The RMSD between the 7 UII models generated by MD (M1-M7) and 5 UII models generated by PEP-FOLD (M1-M5), used as input to PEP-Cyclizer.**

| | | MD M1 | MD M2 | MD M3 | MD M4 | MD M5 | MD M6 | MD M7 | MD M8 | PEP-FOLD M1 | PEP-FOLD M2 | PEP-FOLD M3 | PEP-FOLD M4 |
|---|---|---|---|---|---|---|---|---|---|---|---|---|---|
| MD | **M2** | 1.07 | | | | | | | | | | | |
| | **M3** | 1.10 | 0.93 | | | | | | | | | | |
| | **M4** | 3.20 | 3.34 | 3.15 | | | | | | | | | |
| | **M5** | 2.64 | 2.91 | 2.59 | 1.04 | | | | | | | | |
| | **M6** | 2.23 | 2.52 | 2.25 | 1.45 | 0.77 | | | | | | | |
| | **M7** | 2.49 | 2.70 | 2.43 | 1.38 | 1.00 | 1.15 | | | | | | |
| | **M8** | 2.53 | 2.81 | 2.50 | 1.70 | 1.22 | 1.26 | 0.83 | | | | | |
| PEP-FOLD | **M1** | 1.98 | 2.34 | 2.03 | 1.96 | 1.36 | 1.16 | 1.45 | 1.69 | | | | |
| | **M2** | 1.92 | 2.18 | 2.16 | 3.42 | 2.85 | 2.53 | 2.93 | 2.86 | 2.47 | | | |
| | **M3** | 1.79 | 2.16 | 1.97 | 2.98 | 2.27 | 2.00 | 2.49 | 2.38 | 1.87 | 1.70 | | |
| | **M4** | 2.32 | 2.52 | 2.64 | 3.44 | 3.11 | 2.78 | 2.65 | 2.73 | 2.79 | 2.67 | 2.99 | |
| | **M5** | 2.07 | 2.51 | 2.49 | 4.12 | 3.44 | 3.09 | 3.35 | 3.36 | 2.97 | 2.12 | 2.30 | 2.50 |

| Compound | Code | Sequence | Formula | $MH^{+}calc.^{a}$ | $MH^{+}prat.^{b}$ |
|---|---|---|---|---|---|
| hU-II | LV-4001 | ETPDCFWKYCV | $C_{64}H_{85}N_{13}O_{18}S_{2}$ | 1388.56 | 1388.40 |
| cyclo-[GAG]hUII | LV-4130 | (GAGETPDCFWKYCV) | $C_{71}H_{94}N_{16}O_{20}S_{2}$ | 1555.63 | 1555.63 |
| cyclo-[AGG]hUII | LV-4131 | (AGGETPDCFWKYCV) | $C_{71}H_{94}N_{16}O_{20}S_{2}$ | 1555.63 | 1555.64 |
| cyclo-[GGA]hUII | LV-4132 | (GGAETPDCFWKYCV) | $C_{71}H_{94}N_{16}O_{20}S_{2}$ | 1555.63 | 1555.65 |
| cyclo-[GPA]hUII | LV-4133 | (GPAETPDCFWKYCV) | $C_{74}H_{98}N_{16}O_{20}S_{2}$ | 1595.66 | 1595.66 |
| Nrf2(76-85) | LV-5554 | LDEETGEFL | $C_{46}H_{69}N_{9}O_{19}$ | 1052.47 | 1052.47 |
| Cyclo-[PAA]Nrf2(76-85) | LV-5562 | (PAALDEETGEFL) | $C_{57}H_{84}N_{12}O_{21}$ | 1273.59 | 1273.58 |
| Cyclo-[AGG]Nrf2(76-85) | LV-5565 | (AGGLDEETGEFL) | $C_{53}H_{78}N_{12}O_{21}$ | 1219.54 | 1219.63 |

Table S7: **Peptide characterization**

$^{a}CalculatedMH^{+}$. $^{b}ObservedMH^{+}$ measured by MALDI-ToF mass spectrometry.

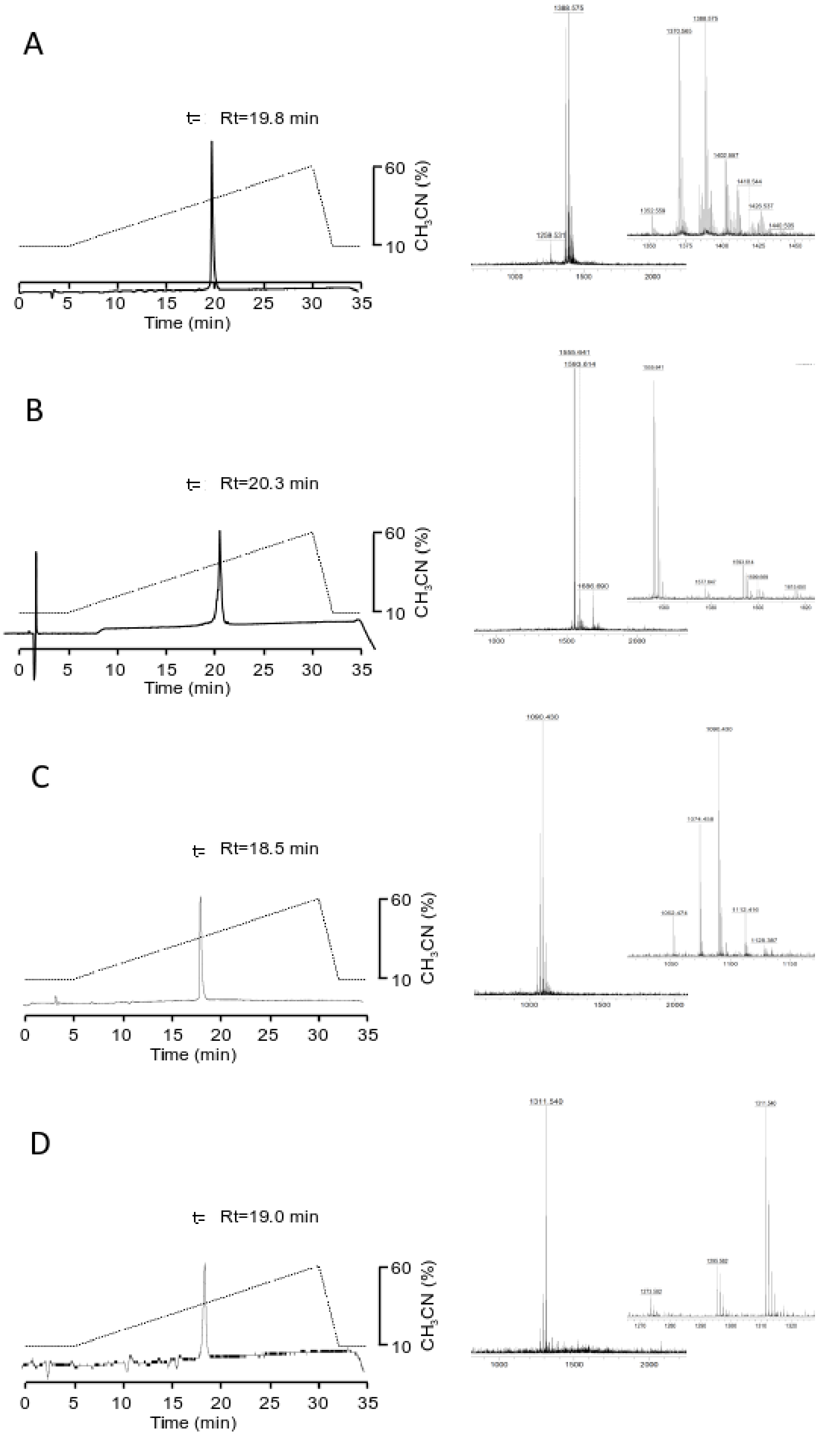


Figure S1: **MALDI-Tof MS and RP-HPLC analysis of synthesized peptides.** A: UII - LV-4001, B: cyclo[AGG]UII - LV-4131, C: Nrf2(76-85) - LV-5554, D: cyclo[PAA]Nrf2(76-85)- LV-5562.

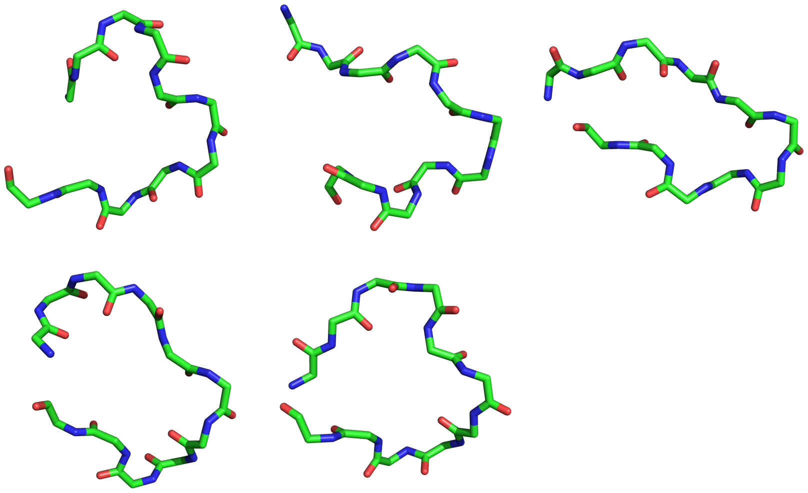

Figure S2: **UII models generated by PEP-FOLD.** The set of 5 UII models generated by PEP-FOLD and refined by MD simulations are shown here.
